# Right-lateralized fronto-parietal network and phasic alertness in healthy aging

**DOI:** 10.1101/826008

**Authors:** Marleen Haupt, Adriana L. Ruiz-Rizzo, Christian Sorg, Kathrin Finke

## Abstract

Phasic alerting cues temporarily increase the brain’s arousal state. In younger and older participants, visual processing speed in a whole report task, estimated based on the theory of visual attention (TVA), is increased in cue compared to no-cue conditions. The present study assessed whether older participants’ ability to profit from warning cues is related to iFC in the cingulo-opercular and/or right fronto-parietal network. We obtained resting-state functional magnetic resonance imaging (rs-fMRI) data from 31 older participants. By combining an independent component analysis and dual regression, we investigated iFC in both networks. A voxel-wise multiple regression in older participants revealed that higher phasic alerting effects on visual processing speed were significantly associated with lower right fronto-parietal network iFC. We then compared healthy older participants to a previously reported sample of healthy younger participants to assess whether behaviour-iFC relationships are age group specific. The comparison revealed that the association between phasic alerting and cingulo-opercular network iFC is significantly lower in older than in younger adults. Additionally, it yielded a stronger association between phasic alerting and right fronto-parietal network iFC in older versus younger participants. The results support a particular role of the right fronto-parietal network in maintaining phasic alerting capabilities in aging.

## Introduction

Warning cues induce short-lived changes in the brain’s “state of readiness” defined as phasic alertness ^1^. In verbal whole report paradigms of briefly presented letter arrays combined with modelling based on the computational Theory of Visual Attention (TVA), previous studies demonstrated that visual ^2^ and auditory warning cues ^3^ increase visual processing speed. Furthermore, we demonstrated that auditory alerting was effective in both healthy younger and older participants ^4^.

Concerning neural correlates, in younger adults, we found that the individual phasic alerting effect on visual processing speed was related to intrinsic functional connectivity (iFC) in the cingulo-opercular network ^5^. IFC was measured in a separate resting-state fMRI session, which allowed to measure ongoing fluctuations of the blood oxygen level dependent (BOLD) signal with a frequency around 0.01-0.1Hz ^6^. Spatial patterns of coherent, i.e. correlated, BOLD fluctuations over time (iFC) constitute intrinsic brain networks which are stable on an intra- ^7^ and inter-subject ^8,9^ level. The cingulo-opercular network (CON) is also referred to as “salience network” ^10^ or “ventral attention network” ^11^. It comprises the anterior prefrontal cortex, anterior insula, frontal operculum, dorsal anterior cingulate cortex, medial superior frontal cortex, and thalamus ^12^.

With respect to older adults, it is unclear whether their preserved enhancement of visual processing speed following alertness cues relies on the same or a different intrinsic brain network. In a comprehensive review on the role of the right-hemisphere in cognitive reserve ^13^, Robertson suggested that the integrity and functional connectivity of the right fronto-parietal network is decisive for the late-life maintenance of attentional abilities, and especially alertness functions ^14^. He summarized evidence for a close, bidirectional relationship of the availability of noradrenaline provided by the locus coeruleus, which is decisive for the ability to increase arousal ^15,16^, and the integrity of the right fronto-parietal network in aging individuals. The model is supported by a more recent demonstration that higher activity of the noradrenergic system (induced by a handgrip task) increased functional connectivity between the locus coeruleus and the fronto-parietal network to a bigger extent for older compared to younger women ^17,18^. Taken together, previous evidence and relevant cognitive reserve models suggest that connectivity in the right fronto-parietal network could be essential for preserved alertness functions in healthy older participants.

The present study, therefore, sought out to investigate whether the individual degree of phasic alerting effects is linked to the individual iFC in the cingulo-opercular network and/or right fronto-parietal network in healthy aging. In order to address this question, we, first, examined the relationship between iFC in the cingulo-opercular network as well as right fronto-parietal network and phasic alerting effects in a group of healthy older adults. Second, we investigated iFC-behaviour relationships in task-related sensory networks. Phasic alerting effects could potentially be related to the auditory network as they were induced by auditory cues and/or visual networks as we measured the effect of these auditory cues on visual processing. In addition to these intra-network analyses, we explored inter-network connectivity pattern between the cingulo-opercular and right fronto-parietal network on the one hand, and the visual and auditory networks on the other hand. In order to ensure the specificity of observed associations, we carried out intra-network and inter-network control analyses in other attention-relevant intrinsic brain networks, expecting that they would not be significantly related to phasic alerting effects. Lastly, we compared iFC-behaviour relationships in the cingulo-opercular and right fronto-parietal network between younger and older healthy adults in order to determine whether associations between phasic alerting effects and iFC are age group specific. In this age group comparison, we contrasted the healthy older participants with a previously reported sample of healthy younger participants ^5^. For all analyses, phasic alerting effects were measured by cue-induced changes in visual processing speed *C* in an offline administered TVA-based whole report paradigm.

## Results

### Phasic alerting effects in healthy older participants

The behavioural data of the original sample of 32 healthy older participants have already been analysed and reported elsewhere ^4^. As one older participant had to be excluded from the present fMRI study due to extensive head motion, we report the behavioural results for 31 healthy older participants included in the present study.

A robust method for the 2×2 repeated-measures design revealed a significant main effect of cueing (*Qa*=6.333, p=.012). Visual processing speed *C* was significantly higher in the cue compared to the no-cue condition. The main effect of CTOA (*Qb*=1.899, p=.168) and the cueing x CTOA interaction (*Qab*=0.046, p=.830) were not significant.

Regarding the main effect of cueing, the Bayes factor of B_10_=1.016 indicates anecdotal evidence for the alternative hypothesis. The main effect of CTOA yields a Bayes factor of B_10_=0.406 (anecdotal evidence for null hypothesis). The cueing x CTOA interaction (B_10_=0.247) demonstrates substantial evidence for the null hypothesis.

In sum, the analyses yielded a significant main effect of cueing and demonstrated that the phasic alerting effects were comparable across the short and long CTOA spectrum in older participants.

### Associations between phasic alerting effects and iFC in healthy older participants

#### Cingulo-opercular and right fronto-parietal network

The cingulo-opercular network encompasses the cerebellum, amygdala, insula, basal ganglia, thalamus, paracingulate gyrus, anterior cingulate cortex, orbital gyrus and frontal gyri (inferior, middle and superior) (see Fig. 1a). The right fronto-parietal networks comprises the cerebellum, prefrontal cortex, frontal gyri, intra-parietal sulcus, inferior parietal lobule, posterior cingulate cortex and temporal gyri (middle and superior) (see Fig. 1b).

**Figure 1.**
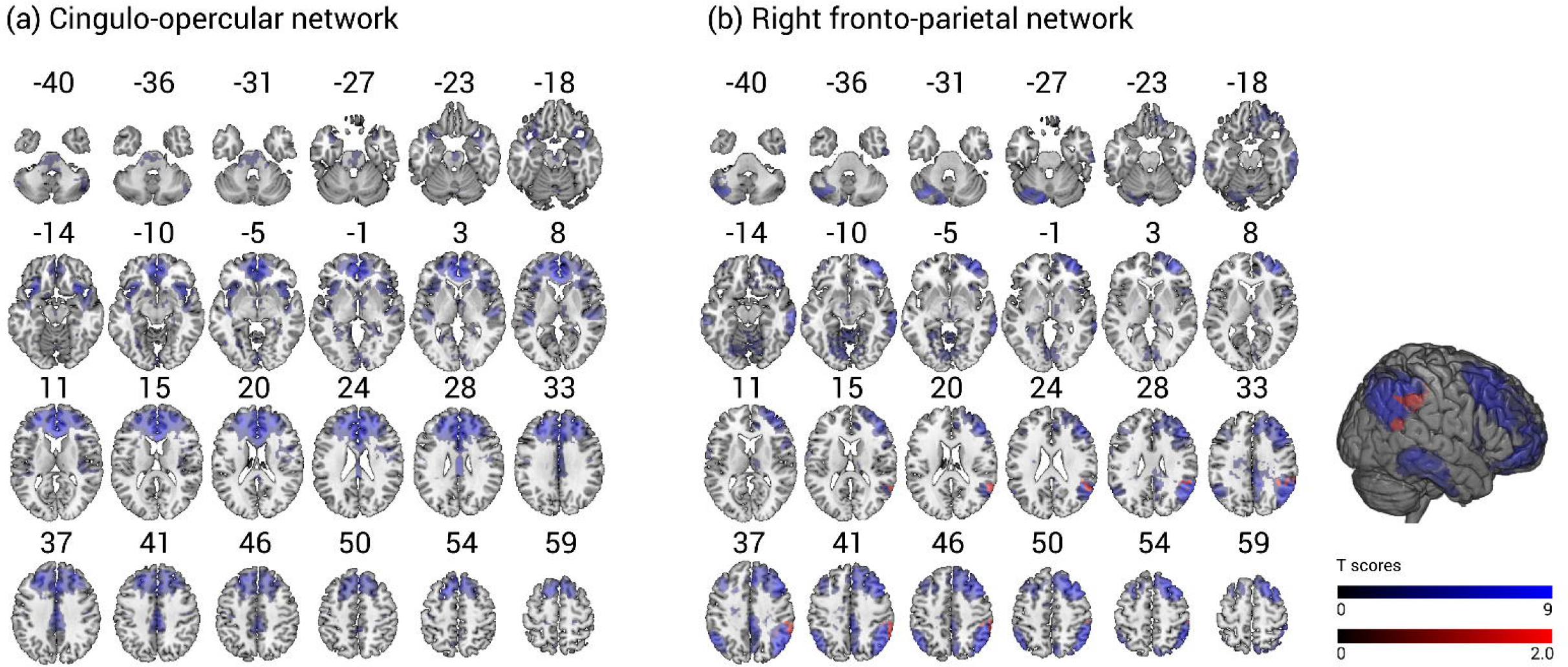
IFC in the cingulo-opercular network (a) and right fronto-parietal network (b) in older healthy participants. Statistical Parametric Mapping of voxel-wise multiple regression of phasic alerting effect on visual processing speed (red) is overlaid on intra-network iFC (blue). The results are obtained by independent component analysis of resting-state fMRI data and are overlaid onto standard anatomical MNI152 templates using the software MRIcroGL (available at: https://www.mccauslandcenter.sc.edu/mricrogl/source); slice numbers in transverse plane are indicated. The results of the multiple regression are controlled for age, sex, head motion, and education (p < .05 FWE corrected at cluster level).

The voxel-wise multiple regression analysis in the group of healthy older participants yielded a significant relationship between the individual degree of phasic alerting effects on visual processing speed *C* and iFC in the right fronto-parietal network (when controlling for age, sex, education, and head motion). Specifically, higher absolute cueing effects averaged over both CTOAs were significantly associated with lower individual iFC in the right fronto-parietal network, peaking in the right superior temporal gyrus (MNI coordinates in mm: [54 - 52 22], cluster size: 647 voxels, T= 4.11, Z=3.56, p=.022, FWE cluster-corrected) (see Fig. 1b). We did not find significant associations between phasic alerting effects and iFC in the cingulo-opercular network.

Importantly, our result should not be interpreted as indicating that the peak region of a significant behaviour-iFC association being particularly “responsible” for phasic alerting effects. As we employed a voxel-wise approach, the iFC values of the significant voxels in the clusters (Z scores) are in principle relative values. This implies that these values can only be interpreted in relation to iFC of other voxels in the corresponding intrinsic brain network. Hence, we can only summarise that the relationship between phasic alerting effects on visual processing speed and iFC in a certain network was best represented in the significant cluster.

#### Auditory and visual networks

In order to address whether iFC in auditory and visual networks is related to phasic alerting effects on visual processing speed following auditory cues, we ran additional multiple regression analyses in these networks (see Supplementary Fig. 1-3). The analyses yielded significant negative associations of phasic alerting effects and iFC in the auditory network with a peak in the left middle frontal gyrus (MNI coordinates in mm: [−28 12 48], cluster size: 521 voxels, T= 4.32, Z=3.70, p=.042 FWE cluster-corrected) and visual networks. For visual network I (Allen component IC39) the peak was located in the left fusiform gyrus (MNI coordinates in mm: [−12 −82 −16], cluster size: 1434 voxels, T= 4.46, Z=3.79, p<.001 FWE cluster-corrected); for visual network II (Allen component IC46) the peak was located in the left lingual gyrus (MNI coordinates in mm: [−30 −58 8], cluster size: 2135 voxels, T= 5.99, Z=4.67, p<.001 FWE cluster-corrected).

#### Control analyses in other attention-relevant networks

To determine whether phasic alerting effects on visual processing speed are specifically linked to iFC in the right fronto-parietal network, we controlled for phasic alerting associations with iFC in other attention-relevant networks. The analyses yielded no significant association of the individual degree of phasic alerting effects on visual processing speed with iFC in the executive control and left fronto-parietal network (all p>.05 FWE cluster-corrected).

#### Inter-network connectivity analyses in healthy older participants

We set out to trace whether the inter-network connectivity of either the cingulo-opercular or the right fronto-parietal networks with other networks is also significantly associated with phasic alerting effects. We entered the above-mentioned attention-related, auditory, and visual networks into an inter-network analysis. For the cingulo-opercular network, the analysis only yielded significant negative correlations with both visual networks. The right fronto-parietal network was positively correlated with the left fronto-parietal, executive control, and auditory networks. The right fronto-parietal network was significantly negatively correlated with both visual networks and did not show any significant correlation with the cingulo-opercular network (see Supplementary Fig. 4).

Importantly, correlation analyses revealed that none of the described significant inter-network correlations were, in turn, significantly correlated with phasic alerting effects (−.229 ≤ all r ≤ .259, all p ≥.159). These results indicate that the decisive link between the phasic alerting effect on visual processing speed *C* and the right fronto-parietal network in older participants is its intra-network iFC and not its inter-network connectivity with other attention-relevant, auditory, or visual networks.

#### Age group comparison of associations between phasic alerting effects and iFC

First, we ran a voxel-wise multiple regression analysis comparing behaviour-iFC relationships in healthy older and younger participants in the cingulo-opercular network. This analysis revealed that the association between the individual degree of phasic alerting effects on visual processing speed *C* and iFC in the cingulo-opercular network (peak located in the left superior orbito-frontal gyrus, MNI coordinates in mm: [−10 30 42], cluster size: 1061 voxels, T= 4.20, Z=3.90, p<.05 FWE cluster-corrected) is significantly lower in older than in younger adults (see Figure 2a).

**Figure 2.**
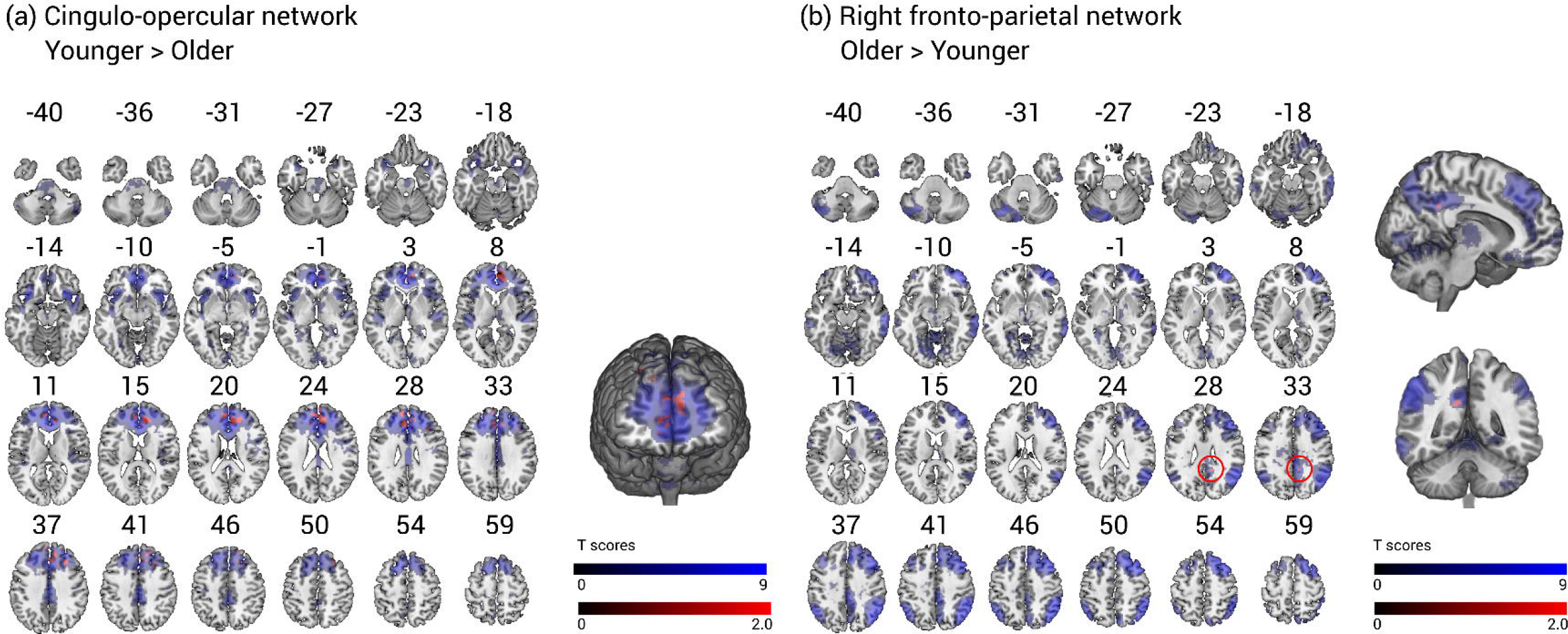
Age group comparison: Voxel-wise multiple regression of phasic alerting effect on visual processing speed split by agegroup is overlaid on iFC (blue) in the cingulo-opercular (a) and right fronto-parietal network (b). The association between phasic alerting effects on visual processing speed and iFC in the cingulo-opercular network is significantly lower in older than in younger adults (a). The association between phasic alerting effects on visual processing speed and iFC in the right fronto-parietal network is higher in older than in younger adults (b). All results are obtained by independent component analysis of restingstate fMRI data and are overlaid onto standard anatomical MNI152 templates using the software MRIcroGL (available at: https://www.mccauslandcenter.sc.edu/mricrogl/source); slice numbers in transverse plane are indicated. All results are controlled for sex, head motion, and education. The red overlays in the cingulo-opercular network are p < .05 FWE corrected at cluster level while the overlays in the right fronto-parietal network are the results of an exploratory analysis (p< .05 uncorrrected at cluster level).

Second, we ran the same analysis in the right fronto-parietal network. The analysis in the right fronto-parietal network did not yield significant differences between age groups in their behaviour-iFC relationship when p < .05 FWE cluster-corrected.

Interestingly, the preceding analysis in only healthy older participants provided strong evidence for a primary behaviour-iFC association in the right fronto-parietal network. Therefore, we decided to perform an additional exploratory analysis without cluster correction but with a conservative voxel-wise threshold of p < .001. This exploratory multiple regression without cluster-level correction suggests that the association between phasic alerting effects and iFC in the right fronto-parietal network is significantly higher in older than younger participants (see Figure 2b). The peak of the association was located in the right posterior cingulate cortex (MNI coordinates in mm: [10 −46 30], cluster size: 18 voxels, T=3.87, Z=3.62, p=.039 at cluster level, uncorrected). This result suggests that there is a trend for a stronger association between phasic alerting effects and iFC in the right fronto-parietal network in older versus younger participants.

## Discussion

Auditory warning cues increase visual processing speed in healthy younger and older participants ^3,4^. In healthy younger adults, such phasic alerting effects on visual processing speed have been linked to intrinsic functional connectivity (iFC) in the cingulo-opercular network ^5^. We wanted to investigate whether the same or distinct functional connectivity patterns do underlie preserved phasic alerting effects in healthy older participants. Previous studies suggested that connectivity in the right fronto-parietal network could be essential for preserved alertness functions in healthy older participants ^14,17,18^.

The present study sought out to identify the underlying neural network mechanisms of this relevant phasic alerting effect in healthy older participants. We addressed this question by relating individual differences in the degree of phasic alerting effects on visual processing speed measured in an offline administered TVA-based paradigm to those in intrinsic functional connectivity (iFC) in the cingulo-opercular and in the right fronto-parietal network. Firstly, we analysed iFC-behaviour relationships in healthy older adults. Secondly, we compared these iFC-behaviour relationships between healthy younger and older adults in order to determine whether associations between phasic alerting effects and iFC are age group specific.

The voxel-wise multiple regression analysis in healthy older participants revealed a significant relationship between the individual degree of phasic alerting effects on visual processing speed and iFC in the right fronto-parietal network. Higher absolute cueing effects (averaged over both CTOAs) were significantly associated with lower individual iFC in the right fronto-parietal network. In contrast to the results in younger participants ^5^, we did not find significant associations between phasic alerting effects and iFC in the cingulo-opercular network in healthy older participants. Importantly, greater connectivity does not automatically imply better behavioural performance. Previous studies have demonstrated that lower behavioural scores were associated with higher iFC in the brain networks of interest ^49,50^. Based on such evidence, Ferreira and Busatto ^51^ summarised that the assumption “the more, the better” is overly simplistic and might be misleading in the interpretation of findings.

We also addressed iFC in, first, the auditory network as we used auditory cues and, second, in visual networks because we assessed alerting effects on visual processing. The analyses yielded significant negative associations of phasic alerting effects and iFC in the auditory network as well as both visual networks. Inter-network connectivity analyses of both the right fronto-parietal network and cingulo-opercular network with these task-relevant networks did not reveal significant relationships which would demonstrate that internetwork connectivity is relevant for inter-individual differences in phasic alertness. However, as we have found significant behaviour-iFC associations in the auditory as well as two visual networks, we suggest that the right fronto-parietal network in healthy older participants is in permanent, bi-directional exchange of information with sensory networks. On the one hand, the information it receives might be dependent on the efficiency of the auditory system to perceive auditory warning signals. On the other hand, the right fronto-parietal network might heighten the readiness of visual networks in order to accelerate the uptake of visual information. The fact that we did not find significant behaviour-related connectivity pattern between networks might indicate that iFC as a measure is not sensitive enough to capture the effectiveness of uni- or bi-directional communication between these brain areas. Potentially, effective connectivity measures could constitute a promising approach to reveal relevant directed, i.e. effective, connectivity patterns between the right fronto-parietal network and the visual or auditory network ^52–54^.

Control analyses were conducted in order to test whether the right fronto-parietal network plays a specific role for alerting effects in older participants. These analyses revealed that neither the intra-network of other attention-related networks nor their inter-network connectivity with the cingulo-opercular or right fronto-parietal network were found to be significantly related to the individual phasic alerting effect.

The comparison of iFC-behaviour relationships in healthy younger versus older participants revealed that associations between phasic alerting effects and iFC are, indeed, age group specific. While phasic alerting effects in younger participants are primarily associated with iFC in the cingulo-opercular network, this associations appears to be driven by the right fronto-parietal network in healthy older participants.

Taken together, the results of the present study complement a previous resting-state fMRI study reporting age-related increases in iFC in the right fronto-parietal network ^49^. Furthermore, the results are in line with reviews and empirical studies proposing a preferential role of right-hemispheric fronto-parietal structures for noradrenaline-mediated attentional, and particularly alertness processes in aging ^14,17,18^. Robertson ^14^ summarized evidence suggesting that the integrity and functional connectivity in this network may facilitate attentional functions, and particularly alertness, in old age. Enhancing visual processing speed in response to a warning signal depicts a central cognitive ability which can temporarily reduce the detrimental impact of age-related declines of general processing resources. Accordingly, older individuals with a high capability to mobilize additional capacity when faced with external warning signals are able to improve their rate of visual information uptake in critical situations. Older adults with lower phasic alertness capabilities might be characterized by a relatively stable capacity, which is i) reduced due to aging effects and ii) does not flexibly adopt to task demands even when provided with alerting environmental cues. A recent study using a TVA-based verbal report paradigm with auditory alerting cues and measuring cue-related power and phase-locking effects in EEG documented that the variability of phasic alertness effects in older adults is reflected in the neural response. Namely, it was found that only those older participants with a relatively youth-like pattern of phase-locking showed reliable performance benefits ^55^. The results of the present study suggest that these inter-individual differences in the ability to utilize auditory phasic alerting cues to increase visual processing speed are linked to iFC in the right fronto-parietal network. They, thereby, support the specific role of the integrity and connectivity of this network for active perception mechanisms and cognitive reserve in old age.

As the present study suggests that, already in healthy aging, phasic alerting effects rely on age-specific underlying spatial patterns of iFC, the question arises how these patterns change in pathological aging. Pathological aging is characterized by significant reductions of visual processing speed, i.e. in mild cognitive impairment and Alzheimer’s disease ^56–58^. Hence, future studies should set out to investigate the degree to which phasic alerting can improve visual processing speed in pathological aging and relate these behavioural effects to underlying iFC patterns.

The present study has several limitations. First, the cross-sectional design and regression analyses do not allow for inferences being drawn regarding directionalities between phasic alertness, iFC in the cingulo-opercular as well as right fronto-parietal network, and aging. Second, our study lacks a direct link to the locus coeruleus-noradrenaline system as we did not include a specific readout. Neuromelanin-sensitive MRI sequences are a promising tool for quantifying locus coeruleus intergrity ^59,60^. In addition, measuring pupil dilation would provide a window into locus coeruleus activity (for a review see ^18^). Third, we cannot directly compare behaviour-iFC relationships between networks as, based on our voxel-wise analysis approach, we neither have one meaningful value representing iFC on the network-level, nor do we have one cluster-based iFC value for each network.

## Methods

### Participants

Thirty-two older adults (≥ 60 years) participated in the present study. One older participant had to be excluded due to extensive head motion. For the age group comparison, they were contrasted with 32 healthy younger (18-35 years) participants whose rs-fMRI data have been reported in a previous study ^5^. The behavioural data of both younger and older participants have already been reported elsewhere ^4^. The final sample of the present study consisted of 32 younger and 31 older participants (see Table 1). All participants reported normal or corrected-to-normal vision, gave informed consent and were reimbursed for their participation. The study was reviewed and approved by the ethics committees of the Department of Psychology of the Ludwig-Maximilians-Universität München and the Klinikum rechts der Isar of the Technical University Munich.

**Table 1.** Demographics and visual processing speed (C) estimates of all participants.

| Variable | Younger participants | Older participants |
| --- | --- | --- |
|  | N = 32 | N = 31 |
| mean age (SD) | 26.6 (4.7) | 71.1 (4.8) |
| sex (female/male) | 20/12 | 10/21 |
| handedness (right/left) | 27/5 | 29/2 |
| mean education in years (SD) | 12.6 (0.9) | 11.8 (1.8) |
| MWTB verbal intelligence score (SD) | 28.6 (4.6) | 32.7 (2.3) |
| MMSE score (SD) | - | 28.9 (1.1) |
| Days between sessions <sup>1</sup> | 240.1 (294.7) | 255.0 (124.3) |
| Absolute Cueing Effect (over both CTOAs)<sup>2</sup> | - | 1.65 (3.65) |
| Absolute Cueing Effect (long CTOA)<sup>3</sup> | 3.92 (5.91) | 1.69 (6.75) |

*Note.* SD: standard deviation; handedness: assessed by Edinburgh Handedness Inventory; MWTB: Mehrfachwahl-Wortschatz-Intelligenztest, maximum score = 37 points; MMSE, Mini-Mental State Examination, maximum score= 30 points with values < 24 points indicating cognitive impairment; <sup>1</sup>Number of days between behavioural and rs-fMRI session; <sup>2</sup> $((C_{\text{cue}}[\text{long}] - C_{\text{no-cue}}[\text{long}]) + (C_{\text{cue}}[\text{short}] - C_{\text{no-cue}}[\text{short}]))/2$ ; <sup>3</sup> $C_{\text{cue}}[\text{long}] - C_{\text{no-cue}}[\text{long}]$

As in the younger sample, the acquisition of the resting-state functional magnetic resonance imaging (fMRI, approx. 1 hour) and the offline TVA-based behavioural assessment (1-1.5 hours) of the older sample took place on two different days. Participants also completed the Edinburgh Handedness Inventory ^19^, a multiple choice German vocabulary test measuring crystallized intelligence, “Mehrfachwahl-Wortschatz-Intelligenztest” (MWTB) ^20^, and the Mini-Mental State Examination (MMSE) as a screening for cognitive impairments indicative of beginning dementia ^21^. None of the participants had to be excluded based on a cut-off criterion for cognitive impairment, i.e. a score below 27/30 points. Demographic information of both study groups is presented in Table 1.

### TVA-based whole report paradigm with alerting cues

The details of the applied TVA-based whole report procedure have already been reported elsewhere ^4^. In short, TVA is closely related to the biased competition account ^22^ and implies parallel processing of several visual objects competing for selection into a capacity-limited visual short term memory (vSTM) store. The probability that an object gets selected before the store is filled is proportional to its processing rate ^23^. An increase of phasic alertness leads to a proportional increase in the processing rate of the object ^24^. The sum of the processing rates of all objects present in the visual display is defined as the observer’s overall visual processing speed *C* (in elements per second) ^25^. By definition, all items in a whole report paradigm share the same expectancy and subjective importance. An increase in the observer’s alertness induced by auditory warning cues will lead to a proportional increase in parameter *C* ^24^. Thus, by comparing visual processing speed in conditions with and without warning cues, the individual phasic alerting effect can be estimated.

All possible trial sequences can be seen in Figure 3a. Participants were instructed to maintain central fixation throughout the task and verbally report all letters recognized with “fair certainty” without any importance of speed or order. After entering all reported letters on the keyboard, the experimenter started the next trial with a button press. A scale presenting the individual’s accuracy rating based on all reported letters succeeded every test block. Participants were asked to maintain an accuracy level between 70% and 90% with a deviating score leading to adapted instructions for the next test block. If the participants’ accuracy rating exceeded 90%, they were asked to also name letters that they believed to have recognized without complete certainty. If participants were less than 70% accurate, they were instructed to only report letters recognized with high certainty even if that meant that they would report fewer letters overall.

**Figure 3.**
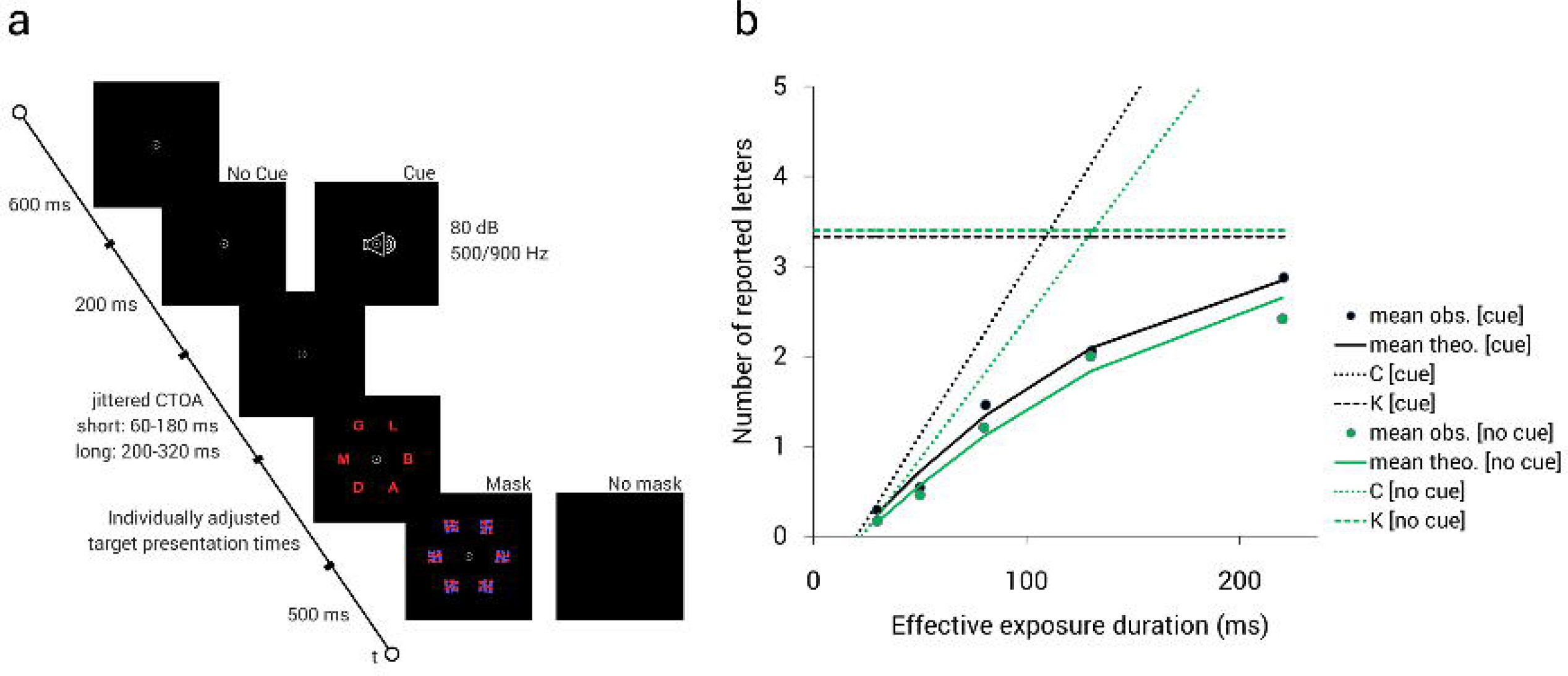
Trial sequence in whole report paradigm (a) and whole report performance for a representative healthy older participant (b). The graph (b) contains a comparison between the cue and no-cue condition. The observed values are displayed as single data points (mean obs.). Solid curves represent the best theoretical fit from the TVA to the observations (mean theo.). The vSTM storage capacity *K* is defined as the asymptote of the function and is marked by a dashed line. The visual perceptual threshold *t0* is defined by the origin of the function (coordinate: t0,0). Visual processing speed *C* is defined as the slope of the function in t0 and is represented by a dotted line.

The whole behavioural experiment consisted of 8 blocks with 84 trials each with half of the trials being preceded by an auditory cue and the other half of trials being uncued ^4^. For the cue as well as no-cue conditions, 2 different cue target onset asynchrony (CTOA) spectrums, “long” and “short,” were used. The “short” CTOAs had an average of 120 ms and were jittered around the average value in steps of ± 20ms, 40ms, and 60ms resulting in an overall range of 60-180ms. The “long” CTOAs had an average of 260ms with the same jittering steps leading to a range of 200-320ms. For each trial, one CTOA was randomly drawn from the according CTOA distribution (short or long). In all conditions, the jittering was balanced with all jittered CTOAs having the same probability and appearing equally often across trials. The effects induced by the auditory alerting cue were comparable across the short and long CTOA spectrums in older participants ^4^. Therefore, we used the average absolute cueing effect calculated as ((C_cue_ [long] – C_no-cue_ [long]) + (C_cue_ [short] – _Cno-cue_ [short]))/2 as the behavioural variable of interest in the subsequent iFC analyses in healthy older participants. Importantly, high-performing younger individuals demonstrated ceiling effects of visual processing speed in the short CTOA, derogating phasic alerting effects. As both age groups demonstrated stable phasic alerting effects in the long CTOA, we only analyzed the absolute cueing effect derived from long CTOA trials in the age group comparison (see Table 1).

The whole report task allows for the estimation of visual processing speed *C*, vSTM storage capacity *K*, and visual perceptual threshold *t0*. For detailed underlying estimation algorithms please refer to Kyllingsbaek ^26^. Figure 2b shows the mathematically modeled exponential growth function of a representative older participant which is relating report accuracy (mean number of reported items) to effective exposure durations. The present study addresses the specific relationship between visual processing speed *C* and iFC as alertness cues predominantly affect visual processing speed ^2–4^.

### Resting-state fMRI

The previously described resting state fMRI data of the younger participants ^5^ are added to the present study to address potential differences between older and younger participants. The acquisition, preprocessing, and analyses of resting-state fMRI data follow the pipeline described in the previous publication.

### Imaging Data Acquisition

Imaging data were acquired on a 3T MR scanner (Philips Ingenia, Netherlands) using a 32-channel SENSE head coil. Small cushions stabilized participants’ heads in the head coil to reduce head motion. Earplugs and headphones reduced scanner noise. Functional data acquisition lasted for 12.5 minutes and participants were instructed to keep their eyes closed, intend to stay awake, and to refrain from performing any cognitive or motor activity, i.e. be at rest, throughout the whole sequence. At the end of the sequence, all participants reported that they had stayed awake. The functional data set consisting of 600 volumes was acquired by multi-band echo-planar imaging (EPI; Preibisch et al., 2015) with a multi-band SENSE acceleration factor of 2 (TR = 1250ms; TE = 30ms; phase encoding in anterior-posterior direction; flip angle = 70°; field of view (FOV) = 192mm^2^; matrix size = 64 × 64, 40 slices with 3mm thickness and an inter-slice gap of 0.3mm; reconstructed voxel size = 3mm × 3mm × 3.29mm). Structural data were obtained by a T1-weighted magnetization-prepared rapid-acquisition gradient echo (MPRAGE) sequence (TR = 9ms; TE = 4ms; flip angle = 8°; FOV = 240mm^2^; matrix = 240 × 240, 170 sagital slices; reconstructed isotropic voxel size=1mm).

### Imaging Data Preprocessing

The resting-state fMRI data were preprocessed in MATLAB (R2017b, version 9.3.0.713579; The Mathworks Inc.) using SPM 12 version 6225 (https://www.fil.ion.ucl.ac.uk/spm/software/spm12/) and the Data Processing Assistant for Resting-State fMRI toolbox version 2.3 (DPARSF) ^28^. After removing the first five functional volumes of every data set to account for T1 saturation effects, slice timing and head motion correction were performed by calling SPM functions. One older participant had to be excluded due to excessive head motion by the criterion of cumulative translation or rotation of 3mm or 3° or due to more than 20% frame-wise displacements >0.5 mm (382/594 displaced frames) ^29^. All images were manually reoriented to the AC-PC axis. The functional images were normalized into Montreal Neurological Institute (MNI) space with a 2-mm isotropic voxel size by unified segmentation to the structural image ^30^. DPARSF integrates the three underlying procedures - coregistration, segmentation (grey matter, white matter, cerebrospinal fluid) and writing normalization parameters - into one processing step. The normalized images were smoothed using a 4mm full-width-at-half-maximum (FWHM) Gaussian kernel. Additionally, band-pass filtering (0.01-0.1Hz) was performed and the effects of nuisance covariates (whole-brain, white matter, and cerebrospinal fluid signals, as well as 12 head motion parameters, their derivatives, and scrubbing regressors) were removed.

### Independent Component Analysis and Dual Regression Analyses

After preprocessing the functional data, we conducted a probabilistic Independent Component Analysis (ICA) in FSL (version 5.0.9) using the MELODIC command-line program version 3.14 ^31,32^. We specified 30 independent components aiming at decomposing the data of the rather heterogeneous sample of healthy younger and older adults into larger networks. We refrained from using more components in order to avoid a split-up of the data into smaller subnetworks. The ICA decomposed each time × space matrix into pairs of time courses and spatial maps on the group level. Subsequently, these files were used as input and a dual regression was employed in order to estimate spatial maps and time courses for each participant ^33,34^. The dual regression approach allows to quantify the functional connectivity of each voxel with each spatial map while controlling for all other spatial maps within each participant ^35^. Most importantly, we chose this approach as dual regression analysis is excelling in detecting inter-individual variability in functional connectivity compared to seed-based functional connectivity analysis ^35^. In a first step, the group-average spatial map was regressed into the individual participants’ time × space matrices, resulting in 30 participant-specific time series. In a second step, the group-average time series was regressed into the same matrices, yielding 30 participant-specific spatial maps, i.e. one per independent spatial map on the group-level. The individual spatial maps contained Z-scores of every voxel within the according map. These Z-scores indicated the similarity of a particular voxel’s time course to the time course of the respective component on the group-level while controlling for all other components. Therefore, the voxel-wise Z-scores were used as input for statistical tests to analyse whether the given component derived Z-scores do relate to behavioural variables. Importantly, the results of the statistical analyses are solely related to the specific output of the ICA, i.e. independent components. These components represent intrinsic brain networks, but the precise brain regions included may vary ^35^. In a last step, the randomise permutation-testing tool (5,000 permutations, FWE-corrected p = .05) within the FSL framework yielded one-sample *t*-test or ‘group’ spatial maps^33,34^.

In order to identify typical intrinsic brain networks with our ICA-dual regression approach, we performed a spatial cross-correlation of our 30 independent components with intrinsic brain network templates derived from Allen et al. ^36^using the *fslcc* command of FSL. Accordingly, we identified the component with the strongest correlation coefficient with the “salience network” (component IC55, r= 0.35) and right fronto-parietal network (component IC60, r=0.54) of Allen et al. ^36^ as the cingulo-opercular and right fronto-parietal network in the present study. These cross-correlations as well as a visual inspection of the brain areas included in the networks ensured that we identified the networks of interest for the present study.

### Statistical Analyses

#### Phasic alerting effects in healthy older participants

Due to non-normal distributions of visual processing speed violating the assumptions of general linear models, we applied an equivalent robust model ^37,38^. We used a robust method based on 20% trimmed means for a 2×2 repeated-measures design with the within subject factors cueing (cue vs. no-cue) and CTOA (short vs. long) for visual processing speed. These analyses were performed using the WRS package ^39^ in RStudio version 1.0.136 ^40^.

Apart from orthodox statistics, we also ran the Bayesian counterpart of repeated-measures ANOVAs ^41^ using JASP version 0.8.5.1 ^42^. JASP calculates the Bayes factor which is a measure for the ratio of the likelihoods of two theories. By comparing those likelihood, the Bayes factor allows for a quantification of the evidence for each theory (e.g. null hypothesis and alternative, experimental hypothesis). Hence, if B_10_ is greater than 3 the present data substantially support the alternative hypothesis while values smaller than 1/3 are substantially favour the null hypothesis. B_10_ values between 1 and 3 (as well as 1 and 1/3 accordingly) solely yield anecdotal evidence for an hypothesis ^43,44^.

### Associations between phasic alerting effects and iFC in healthy older participants

#### Cingulo-opercular and right fronto-parietal network

The individual spatial maps resulting from the described second step of the dual regression served as input for the intra-network analyses conducted in SPM12 (http://www.fil.ion.ucl.ac.uk/spm/software/spm12/). We performed two voxel-wise multiple regressions of the absolute cueing effect averaged over both CTOAs on iFC in the cingulo-opercular and right fronto-parietal network. For these analyses and the following associating behavioural measures and iFC within a given network, we performed significance testing for significance threshold p < .05 together with family-wise error correction for multiple comparisons at the cluster level (FWE cluster-corrected). We added age, sex, education, and head motion as planned covariates. We controlled for head motion by adding mean volume-to-volume head motion, i.e. frame-wise displacement, as a covariate to the multiple regression. We chose the measure by Jenkinson et al. ^45^ as it considers voxel-wise differences in its derivation ^46^.

#### Auditory and visual networks

We also identified task-relevant, sensory networks by visual inspection and cross-correlation. Subsequently, we tested for a significant relation between alerting effects and iFC in two visual networks (IC39, r=0.37; IC46, r=043) and an auditory network (IC17, r=0.34) as our behavioural task consisted of visual stimuli and contained an auditory cue.

#### Control analyses in other attention-relevant networks

In order to address the specificity of the relationship between phasic alerting effects and iFC, we additionally performed control analyses in other attention-relevant networks. We identified them by visual inspection and cross-correlation with templates by Allen et al. (2011) as we did with the two networks of interest. We chose to control for alerting associations with iFC in the executive control network (IC71, r=0.47) and the left fronto-parietal network (IC52, r=0.42) as both networks have been reported to be associated with attentional processes in fMRI task studies ^1,47^. The executive control network is distinguishable from the left and right fronto-parietal networks as it contains bilateral temporal gyri, bilateral precuneus, and the right precentral gyrus ^36^. It does neither include frontal structures nor is it restricted to one hemisphere.

#### Inter-network connectivity analyses in healthy older participants

Furthermore, we explored whether phasic alerting effects are significantly associated with inter-network functional connectivity pattern between the cingulo-opercular or right fronto-parietal and task-relevant (auditory and visual) networks. We also controlled for inter-network connectivity between the two networks of interest and other attention-relevant networks. We addressed these questions by entering the individual time courses of the mentioned intrinsic brain networks (yielded by the first step of dual regression) into an inter-network analysis (using custom code written in MATLAB; also see ^48^. We correlated the time course of the cingulo-opercular and right fronto-parietal network with the ones derived from the other five attention- and task-relevant networks of interest per participant. Subsequently, we performed Fisher r-to-Z transformation and correlation analyses to test whether the inter-network connectivity patterns were significantly correlated with the absolute cueing effect averaged over both CTOAs.

### Age group comparison

Finally, to address whether previously analysed associations between phasic alerting effects and iFC in healthy older participants are, indeed, age group specific, we entered healthy older as well as younger participants into intra-network analyses in the cingulo-opercular as well as right fronto-parietal network. For both networks, we performed voxel-wise multiple regressions of the absolute cueing effect in the long CTOA split by age group on iFC values (p < .05 FWE corrected for multiple comparisons at the cluster level). We compared a vector including values for the absolute cueing effect for younger participants and zeros for older participants with a vector containing absolute cueing effect values for older participants and zeros for younger participants. As we were interested in the interactive effect of phasic alerting and age group on iFC, we controlled for the main effects of, both, age group and absolute cueing effect. Additionally, we added sex, education, and head motion as planned covariates.

## Abbreviations

TVA: Theory of Visual Attention of Bundesen
iFC: intrinsic functional connectivity
rs-fMRI: resting-state functional magnetic resonance imaging
vSTM: visual short term memory

## Data availability

The data and code used in the present study are available upon direct request and the data sharing complies with the institutional ethics approval.

## Acknowledgements

We thank Julia Neitzel and Satja Mulej Bratec for their support with the fMRI data acquisition.

This work was supported by the German Forschungsgemeinschaft [grant number FI 1424/2e1 to KF and grant number SO 1336/1e1 to CS]; the European Union’s Seventh Framework Programme for research, technological development and demonstration [INDIREA, grant number ITN-2013-606901 to KF]; and a research scholarship of the Elite Network of Bavaria to MH.

## Author contributions

K.F. and C.S. conceived and designed the study. M.H. led and coordinated the data acquisition. M.H. and A.R.-R. analysed the fMRI data. M.H. analysed the behavioural data. K.F. and C.S. aided in the interpretation of the data. M.H. drafted the main manuscript. All authors repeatedly reviewed the drafts and revised the manuscript critically for important intellectual content.

