## Supplementary for "Right-lateralized fronto-parietal network and phasic alertness in healthy aging"

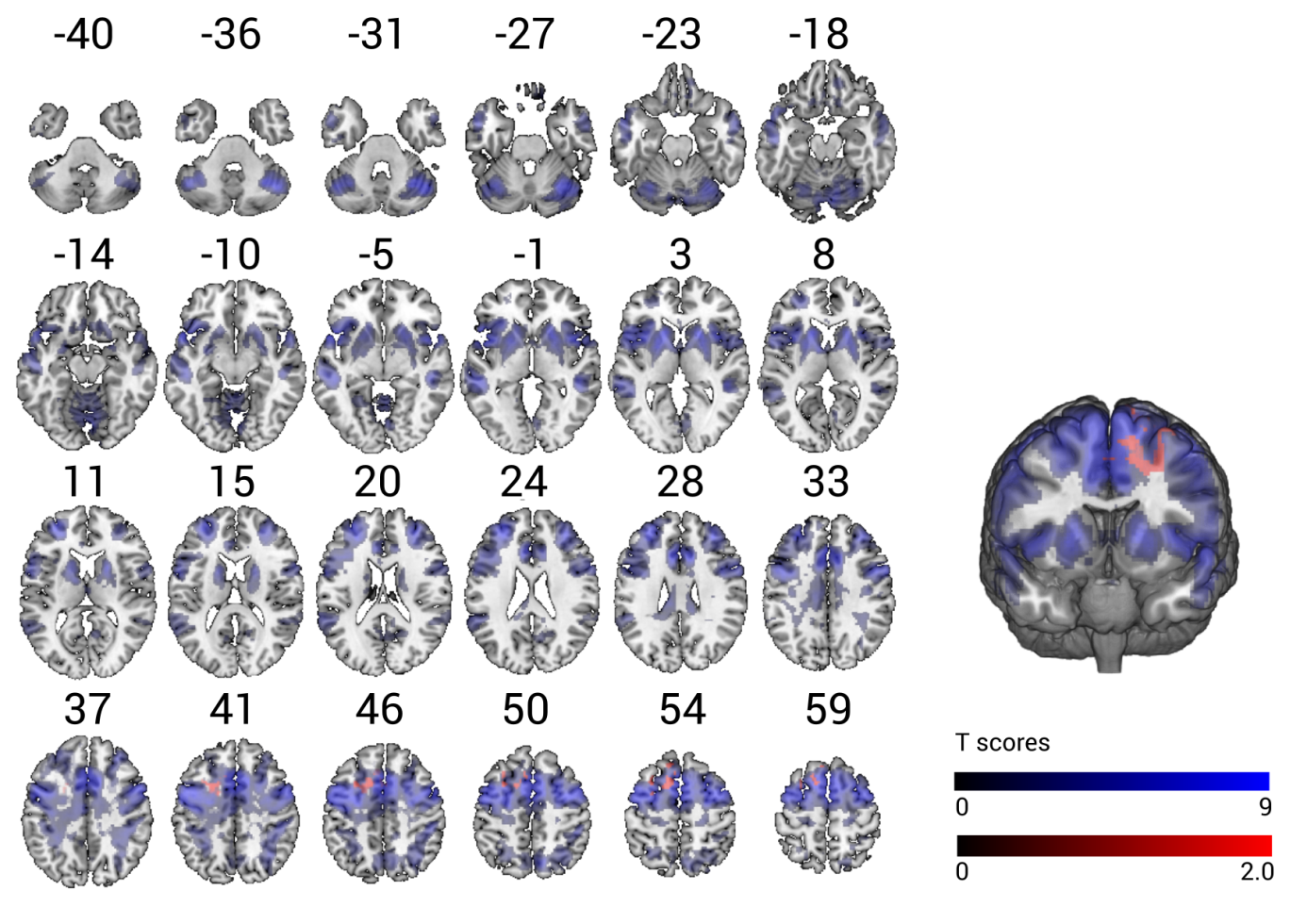

*Figure S1*. Statistical Parametric Mapping of voxel-wise multiple regression of phasic alerting effect on visual processing speed (red) is overlaid on iFC in auditory network (blue). The results are obtained by independent component analysis of resting-state fMRI data and are overlaid onto standard anatomical MNI152 templates using the software MRIcroGL (available at: https://www.mccauslandcenter.sc.edu/mricrogl/source); slice numbers in transverse plane are indicated. The results of the multiple regression are controlled for age, sex, head motion, and education (p < .05 FWE corrected at cluster level).

Marleen Haupt, Adriana L. Ruiz-Rizzo, Christian Sorg, and Kathrin Finke

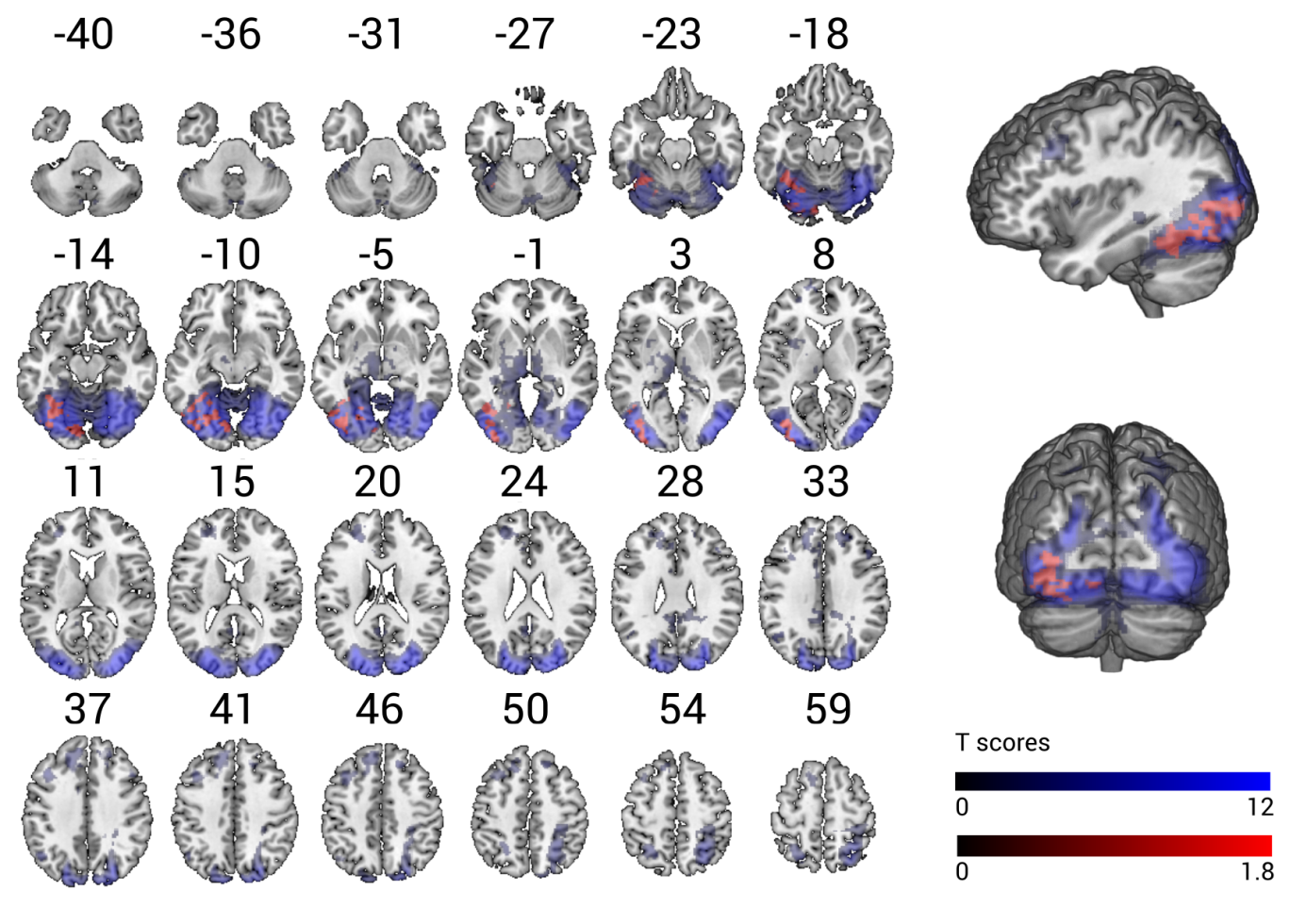

*Figure S2*. Statistical Parametric Mapping of voxel-wise multiple regression of phasic alerting effect on visual processing speed (red) is overlaid on iFC in visual network I (blue). The results are obtained by independent component analysis of resting-state fMRI data and are overlaid onto standard anatomical MNI152 templates using the software MRIcroGL (available at: https://www.mccauslandcenter.sc.edu/mricrogl/source); slice numbers in transverse plane are indicated. The results of the multiple regression are controlled for age, sex, head motion, and education (p < .05 FWE corrected at cluster level).

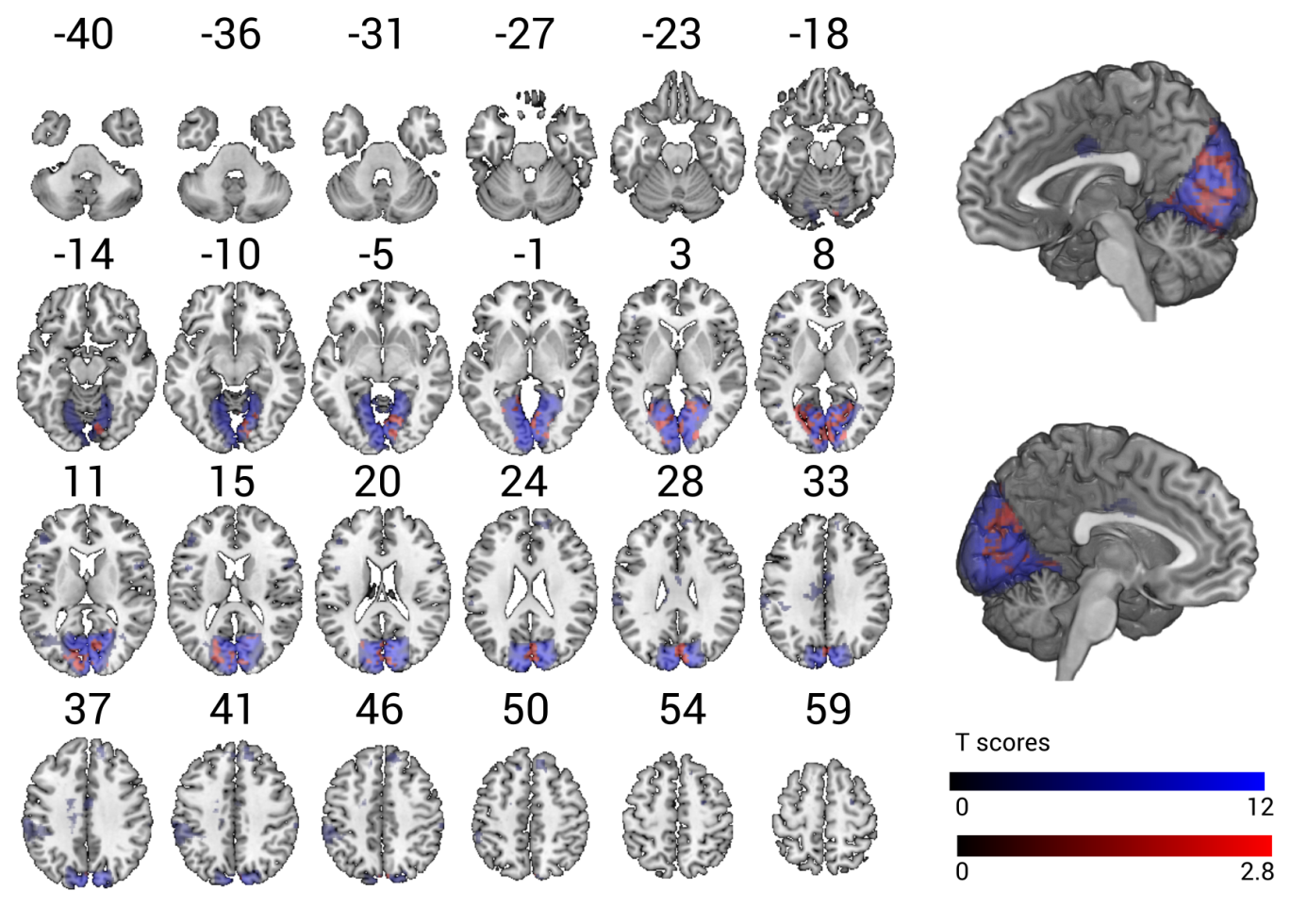

*Figure S3*. Statistical Parametric Mapping of voxel-wise multiple regression of phasic alerting effect on visual processing speed (red) is overlaid on iFC in visual network II (blue). The results are obtained by independent component analysis of resting-state fMRI data and are overlaid onto standard anatomical MNI152 templates using the software MRIcroGL (available at: https://www.mccauslandcenter.sc.edu/mricrogl/source); slice numbers in transverse plane are indicated. The results of the multiple regression are controlled for age, sex, head motion, and education (p < .05 FWE corrected at cluster level).

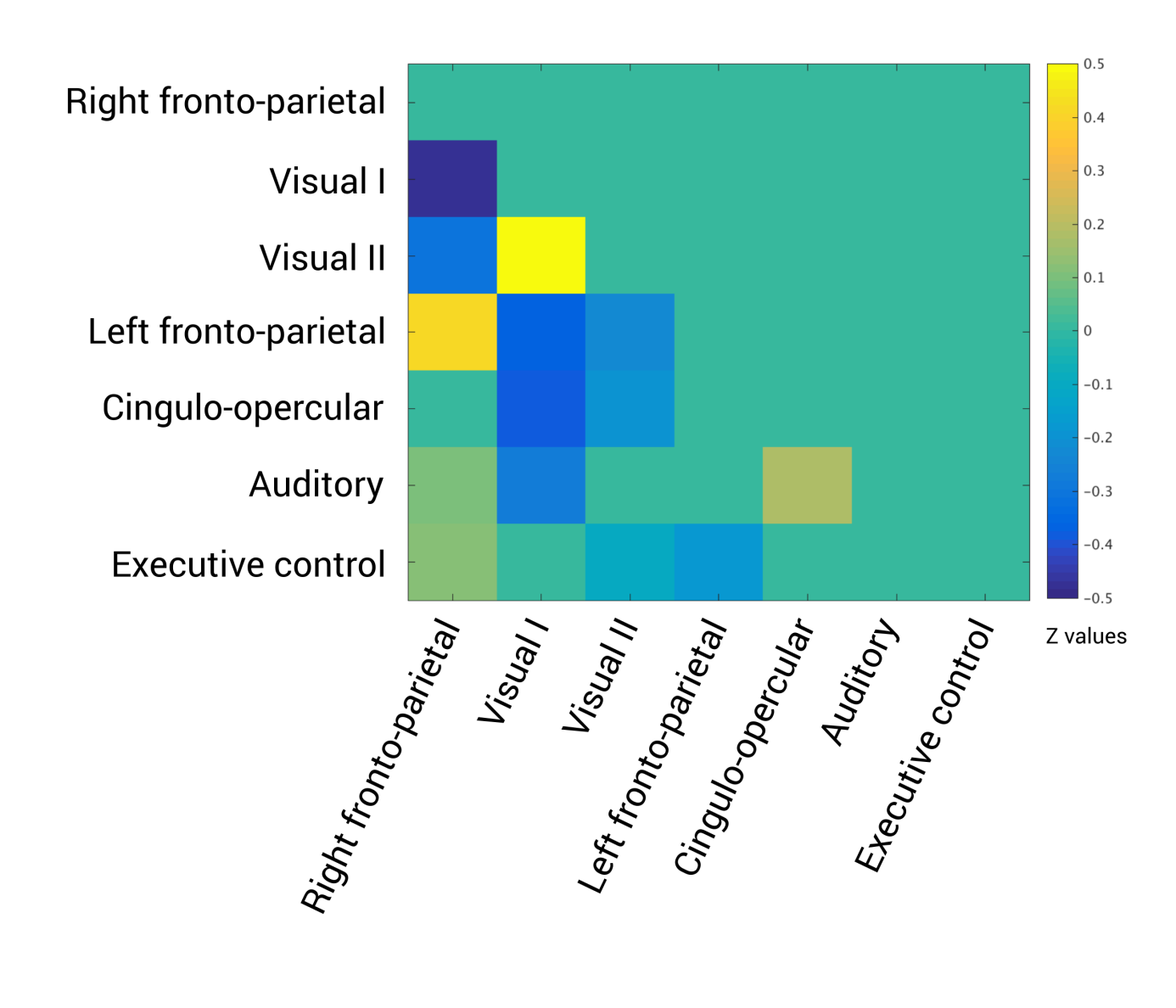

*Figure S4*. Inter-network connectivity of cingulo-opercular and right fronto-parietal network with other attention-related, auditory, and visual networks. The figure displays results of one-sample t-tests (p<.05, FDR corrected for multiple comparisons) of the correlations between networks. Significant positive correlations are highlighted by warm colours, significant negative correlations are reflected by cool colours. The colour bar indicates mean Fisher r-to-z transformed values.
